# The P2X7 receptor localizes to the mitochondria, modulates mitochondrial energy metabolism and enhances physical performance

**DOI:** 10.1101/2020.09.18.303511

**Authors:** Alba Clara Sarti, Valentina Vultaggio-Poma, Simonetta Falzoni, Sonia Missiroli, Anna Lisa Giuliani, Paola Boldrini, Massimo Bonora, Francesco Faita, Nicole Di Lascio, Claudia Kusmic, Anna Solini, Salvatore Novello, Michele Morari, Marco Rossato, Mariusz R. Wieckowski, Carlotta Giorgi, Paolo Pinton, Francesco Di Virgilio

## Abstract

Basal expression of the P2X7 receptor (P2X7R) improves mitochondrial metabolism, ATP synthesis and overall fitness of immune and non-immune cells. We investigated P2X7R contribution to energy metabolism and subcellular localization in fibroblasts (mouse embryo fibroblasts and HEK293 human fibroblasts), mouse microglia (primary brain microglia and the N13 microglia cell line), and heart tissue. The P2X7R localizes to mitochondria, and its lack a) decreases basal respiratory rate, ATP-coupled respiration, maximal uncoupled respiration, resting mitochondrial potential, mitochondrial matrix Ca^2+^ level, b) modifies expression pattern of oxidative phosphorylation (OxPhos) enzymes, and c) severely affects cardiac performance. Hearts from *P2rx7*-deleted versus WT mice are larger, heart mitochondria smaller, and stroke volume (SV), ejection fraction (EF), fractional shortening (FS) and cardiac output (CO), are significantly decreased. Accordingly, physical fitness of P2X7R-null mice is severely reduced. Thus, the P2X7R is a key modulator of mitochondrial energy metabolism and a determinant of physical fitness.

## Introduction

Adenosine 5’-triphosphate (ATP), the universal energy currency, sustains all cellular functions and responses, and at the same time is also an extracellular messenger involved in cell-to-cell communication in virtually every tissue (Burnstock 2014). As an extracellular messenger, ATP is of remarkable importance at sites of cell damage or distress since it is one of the earliest and most important damage-associated molecular patterns (DAMPs) released (Bours et al. 2006, Bours et al. 2011). Due to this dual role, as a DAMP and a high energy intermediate, changes in the extracellular ATP concentration in response to pathogens or to sterile injury might serve not only as alarm signals but also as triggers to activate cellular energy synthesis in conditions of high energy demand. This would make cells more reactive to the impending danger.

Virtually all cells are equipped with plasma membrane receptors for extracellular ATP, the P2 receptors (P2Rs) (Burnstock 2006). P2Rs mediate a multiplicity of responses in the nervous, endocrine, immune and cardiovascular systems. P2Rs are comprised of two subfamilies, the P2Y metabotropic (P2YR) and the P2X ionotropic (P2XR) receptors, each numbering eight and seven members, respectively. Within the P2XR subfamily, the P2X7R subtype is of interest for its peculiar permeability properties and the role in inflammation and cancer (Di Virgilio et al., 2017; Di Virgilio et al., 2018). This receptor is in fact a very potent activator of the NLRP3 inflammasome, of IL-1β processing and release (Di Virgilio 2013), as well as a strong promoter of cancer cell proliferation (Di Virgilio and Adinolfi 2016), and, when over-activated, a trigger of cell death (Di Virgilio et al. 1989). Survival-promoting effects are likely due to P2X7R capacity to support oxidative phosphorylation and glycolysis, and thus enhance intracellular ATP synthesis (Adinolfi et al. 2005, Amoroso et al. 2012). Since during physical activity or at sites of inflammation the extracellular ATP concentration is several fold increased (Forrester 1966, Wilhelm et al. 2010), the P2X7R might be considered a bioenergetics sensor enabling the intracellular ATP synthetic apparatus to sense extracellular ATP levels, and thus meet the increased energy demand. Along these lines, stimulation of the P2X7R has been previously shown to enhance energy metabolism in mice (Giacovazzo et al. 2019). In this study P2X7R subcellular localization and its effect on mitochondrial energy metabolism was investigated. As cellular models, we used cell types well-known for P2X7R expression and function, i.e. fibroblasts (mouse embryo fibroblasts, MEFs, and P2X7R-transfected HEK293, HEK293-P2X7R, human fibroblasts), and microglia (primary brain microglia and the N13 microglia cell line). Mouse fibroblasts and mouse microglia were isolated from WT or *P2rx7*-deleted mice. N13 microglia was available as the WT cell line or the N13 R cell line selected for low P2X7R expression. Finally, to fully appreciate the functional impact of P2X7R-dependent mitochondrial dysfunction in a tissue heavily dependent on oxidative phosphorylation, we investigated heart performance in WT and P2X7R-deleted mice. Altogether, these data show that the P2X7R localizes to the mitochondria in different cell types, and its lack impairs oxidative phosphorylation, affects cardiac performance and decreases physical fitness.

## Results

### Lack of the P2X7R impairs energy metabolism

In a previous study we showed that P2X7R expression in human HEK293 fibroblasts increases mitochondrial Ca^2+^ concentration, mitochondrial membrane potential and overall intracellular ATP content (Adinolfi et al. 2005). **Figure 1A-C** and **Figure 1E-G** show that P2X7R down-modulation or *P2rx7* gene deletion decreases resting mitochondrial potential and severely impairs all respiratory indexes in N13 microglia cells, primary mouse microglia, and mouse embryo fibroblasts (MEFs). Accordingly, overexpression of the P2X7R in cells lacking the endogenous receptor (e.g. human HEK293 cells) increases mitochondrial potential and improves all respiratory indexes (**Figure 1D,H**). Partially depolarized mitochondria are anticipated to have lower matrix Ca^2+^ levels as the driving force for Ca^2+^ is reduced. To verify this prediction we measured mitochondrial Ca^2+^ with the FRET-based GCaMP indicator. As shown in **Figure 1I**, mitochondrial matrix Ca^2+^ level is significantly lower in HEK293 WT versus P2X7R-transfected HEK293 cells (HEK293-P2X7R). Previous data showed that NADH activity is modulated by the mitochondrial matrix Ca^2+^ concentration (Griffiths and Rutter 2009), thus we verified whether lack of P2X7R is associated with changes in the NADH concentration. In HEK293-P2X7R cells, higher matrix Ca^2+^ concentration is associated to lower intra-mitochondrial NADH levels (**Figure J**), and accordingly MEFs from *P2rx7*-deleted mice and mouse microglia selected for low P2X7R expression (N13R microglia) show higher intra-mitochondrial NADH levels (**Figure 1K-M**).

**Figure 1.**
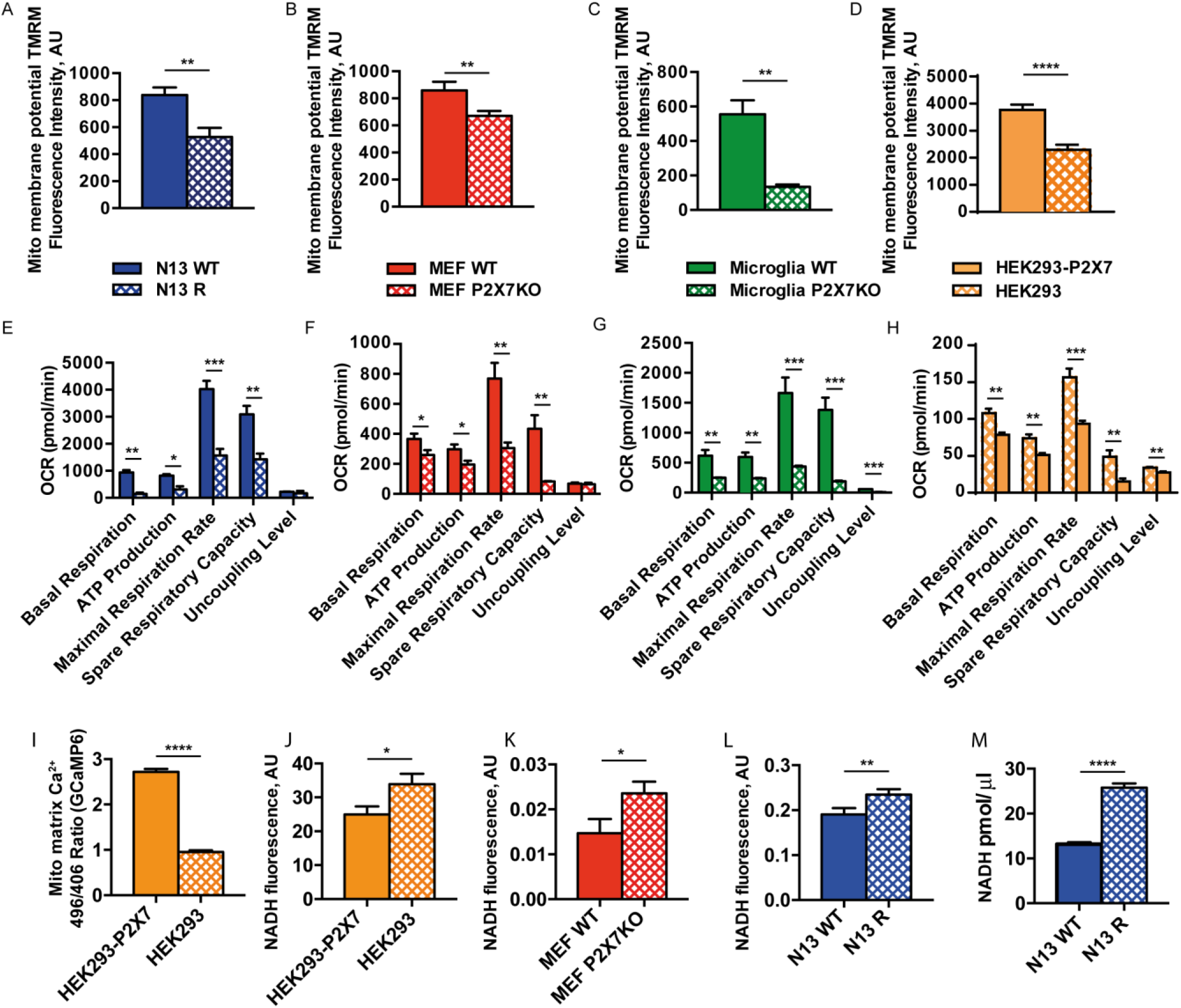
Lack of the P2X7R impairs mitochondrial potential, oxygen consumption, calcium level and causes intra-mitochondrial NADH accumulation. Cells, plated on 12-mm glass coverlips (**A**,**B**,**C**,**D**), were loaded with TMRM (20 nM) at 37°C for 20 min in KRB buffer. Fluorescence was measured with a Zeiss LS510 confocal microscope. Images were then analyzed with ImageJ software. Mitochondrial membrane potential (Ψm) was expressed as the ratio between TMRM fluorescence (in arbitrary units, AU) before and after carbonyl cyanide 4-(trifluoromethoxy) phenylhydrazone (FCCP) addition. Exact fluorescence AU values: N13 WT (average ± SEM = 838.5 ± 55.96, n = 13) or N13 R (average ± SEM = 527.9 ± 66.60, n = 14) (**A**); MEFs WT (average ± SEM = 886.6 ± 59.86; n = 31) or MEFs P2X7-KO (average ± SEM = 671.5 ± 35.49, n = 43) (**B**); primary WT microglia (average ± SEM = 551.1 ± 81.53, n = 20) or primary P2X7-KO microglia (average ± SEM = 134.0 ± 13.69, n = 8) (**C**); HEK293-P2X7 (average ± SEM = 3769 ± 191.0, n = 47) or HEK293 (average ± SEM = 2296 ± 187.7, n = 45) (**D**). Alternatively, cells were plated in XF96 96-well cell culture plate (**E,F,G,H**) and analyzed for oxygen consumption in a SeaHorse apparatus. Exact averages ± SEM for panels **E**, **F**, **G**, **H** are shown in Supplementary Table I. Oxygen consumption rate (OCR) was normalized on cell content. Data are presented as average ± SEM of n = 3 for panels **E**, **F**, **G** and n = 7 for panel **H**. HEK293-P2X7R (average arbitrary fluorescence units ± SEM = 2717 ± 0.069, n = 57) or HEK293 (average arbitrary fluorescence units ± SEM = 0.95 ± 0.039, n = 58) cells were transfected with the mitochondrial-selective FRET-based, GCaMP fluorescent Ca^2+^ indicator (**I**), or analysed for NADH autofluorescence with a fluorescence microscope (average arbitrary fluorescence units ± SEM, HEK293-P2X7R = 23.33 ± 2.07, n = 25; HEK293 = 31.31 ± 2.92, n = 31) (**J**). MEFs WT (average arbitrary fluorescence units ± SEM = 0.015 ± 0.002, n = 11) or MEFs P2X7R-KO (average arbitrary fluorescence units ± SEM = 0.026 ± 0.002; p = 0.006, n = 23) (**K**), and N13 WT (average arbitrary fluorescence units ± SEM = 0.19 ± 0.01, n = 24) or N13 R (average arbitrary fluorescence units ± SEM = 0.23 ±0.01, n = 37) (**L**), were also analysed for NADH fluorescence. Fluorescence emission was acquired with an IX-81 Olympus automated epifluorescence microscope as described in Materials and Methods. Intracellular content (pmol/μL) of NADH for N13 WT (average ± SEM = 13.18 ± 0.25, n = 3) and N13 R (average ± SEM = 25.80 ± 0.39, n = 3) (i) was also measured with a NAD/NADH Assay Kit, as described in Methods. P values are calculated with the two-tailed unpaired Student’s t-test; * p < 0.05; ** p < 0.01; *** p < 0.001; ****, p < 0.0001.

Based on the known Ca^2+^ dependency of mitochondrial dehydrogenases (Griffiths and Rutter 2009), we anticipated that NADH levels should be lower in the presence of lower matrix Ca^2+^ concentrations, while in fact they are higher. Thus, we hypothesized that accumulation of NADH in the absence of P2X7R could be rather due to reduced Complex I activity. To support this hypothesis, we found that HEK293-P2X7R and P2X7R-sufficient N13 WT cells show increased Complex I protein levels compared to HEK293-WT and P2X7R-deficient N13 R cells, respectively (**Figure 2A-D**). It is therefore likely that lower matrix Ca^2+^ in P2X7R-deficient cells is the result of the reduced mitochondrial potential due to lower Complex I expression, and thus to reduced respiration as shown in Figure 1. We also observed reduced Complex II expression in HEK293-P2X7R versus HEK293-WT cells, while no changes in Complex II expression are found in N13 WT versus N13 R cells. We then estimated total mitochondrial content of N13 and MEF cells by measuring expression of the mitochondrial markers TOM20, TIM23 and Hsp60. As shown in Supplementary Information 1, mitochondrial markers are not significantly reduced in the absence of the P2X7R. In N13 cells, reduced expression of the P2X7R decreases mitochondrial membrane potential (**Figure 2G**) and impairs P2X7R physiological responses such as motility (**Figure 2E,F**) and oxygen radical generation (**Figure 2H**). All these responses are restored by supplementation of methyl-succinate, a Site II electron donor that bypasses Site I, and therefore compensates reduced NADH dehydrogenase activity.

**Figure 2.**
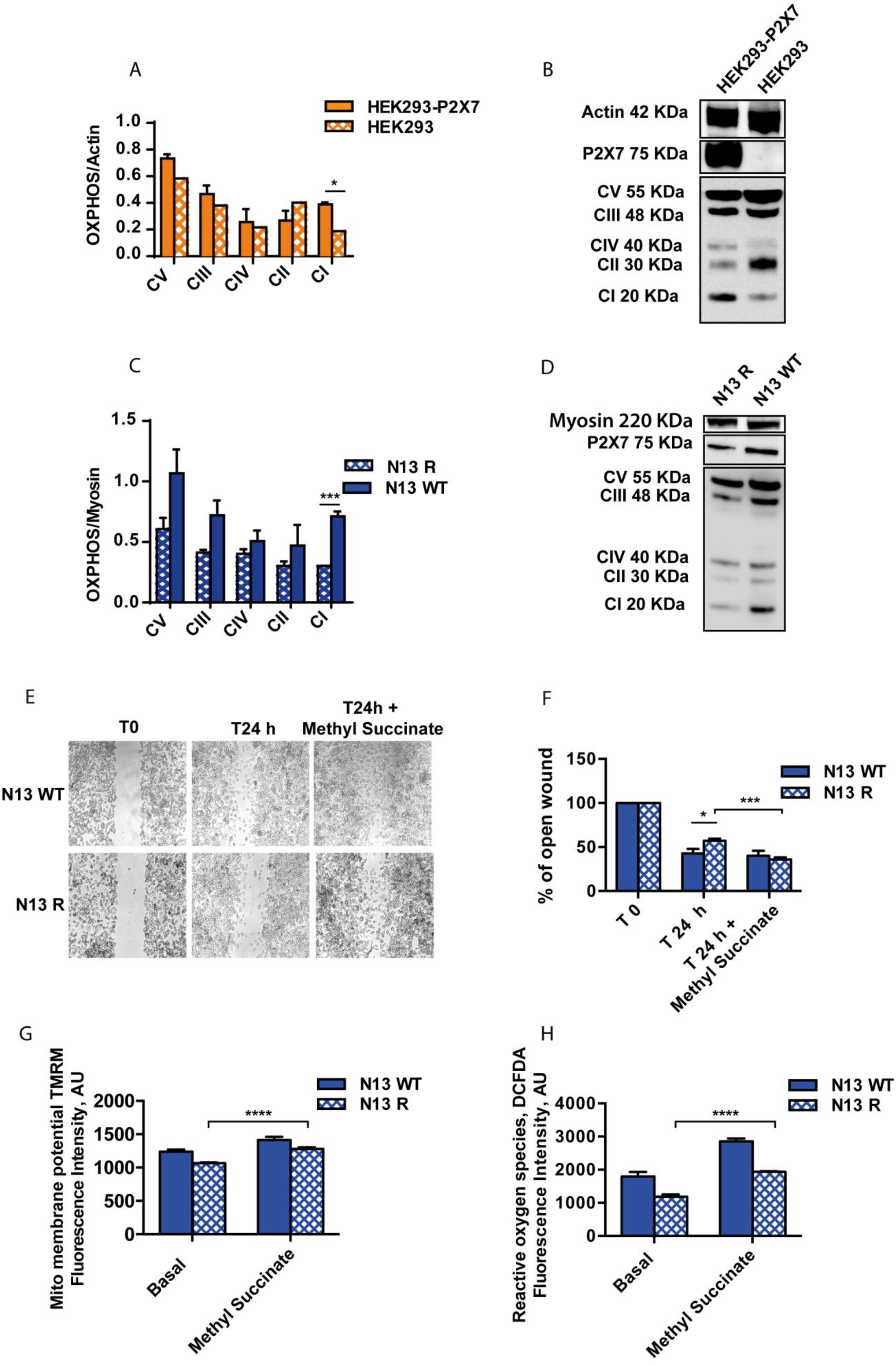
Lack of the P2X7R decreases Complex 1 content of mitochondrial respiratory chain and compromises physiological responses of cells.. Densitometry (**A**,**C**) and Western Blot (**B**,**D**) for respiratory chain complexes (OXPHOS) from HEK293 or HEK293-P2X7R, and N13 WT or N13 R cells is shown. Average absorbance (AU) ± SEM for HEK293-P2X7 = 0.389 ± 0.0141, n = 3; HEK293 = 0.1886 ± 0.059, n = 3; N13 R = 0.301 ± 0.0018, n = 3; N13 WT = 0.712 ± 0.039, n = 3. Ten μg of protein was loaded in each lane. Scratch wound assay in N13 WT or N13 R culture (**E**). Cells were grown in 24-well plates and the wounds were made with a sterilized one-milliliter pipette tip at the same time in all wells at 80% confluence. Wound width (**F**) was measured with Image J at time 0 (T0) and after 24 h (T24). Average ± SEM wound width at T24 as a percentage at T0 was 42.67 ± 5.28, n = 5, for N13 WT, and 57.11 ± 2.269, n = 5, for N13 R. In the presence of 2.5 mM methylsuccinate wound width at T24 was 35.90 ± 2.39 for N13 R, and 40.14 ± 5.64, n = 5, for N13 WT. Mitochondrial potential in the absence or presence of 2.5 mM methyl succinate was measured by TMRM fluorescence with an epifluorescence microscope (average arbitrary fluorescence units ± SEM, N13 R basal 1075 ± 16.19, n = 35, vs N13 R+methylsuccinate 1265 ± 29.35, n = 35) (**G**). Oxygen radical production in the absence or presence of 2.5 mM methyl succinate was measured by 2’,7’-dichlorodihydrofluorescein (DCF) with a Tali image-based cytometer (average fluorescence arbitrary units ± SEM for N13 WT basal 1795 ± 68.67, n = 438; N13 R basal 1184 ± 34.20, n = 347); N13 WT + methylsuccinate 2854 ± 111.7, n = 439; N13 R + methylsuccinate 1933 ± 41.26, n = 383) (**H)**, as described in Materials and Methods. P values are calculated with the two-tailed unpaired Student’s t-test. *, p < 0.05; **, p < 0.01; ***, p < 0.001; ****, p < 0.0001.

### The P2X7R localizes to the mitochondria

Previous data hinted to a P2X7R localization to the nuclear membrane (Atkinson et al. 2002), and possibly to phagosomes (Kuehnel et al. 2009). Due to its dramatic effect on oxidative phosphorylation, we hypothesized that the P2X7R, besides its canonical plasma membrane expression, might also localize to the mitochondria. To address this issue, we probed mitochondrial fractions from cells natively expressing the P2X7R, i.e. MEF WT, and N13 WT, or transfected with the P2X7R, i.e. HEK293-P2X7R, with an anti-P2X7R antibody raised against the C-terminal domain. As shown in **Figure 3A-C**, a band of the approximate MW of 75 kDa is labeled in mitochondrial fractions from the three different cell populations. A band corresponding to the P2X7 subunit is clearly visible not only in a mitochondrial preparation obtained according to standard fractionation procedures, but also in a highly purified mitochondrial fraction (HMF) enriched in mitochondrial calcium uniporter (MCU) content and fully lacking IP3 receptor 3 (IP3R3) and β-tubulin.

**Figure 3.**
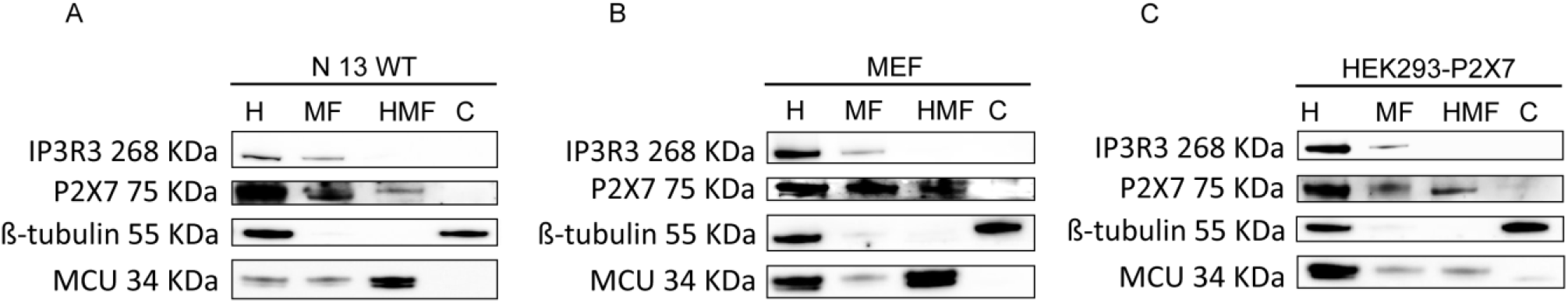
The P2X7R localizes to the mitochondria. Western blot analysis of lysates (10 μg of protein/lane) from whole cells (H), a crude mitochondrial fraction (MF) or a highly purified mitochondrial fraction (HMF). Cytosolic fraction (C) was loaded as control. Cell fractionation was performed as described in Materials and Methods. Inositol phosphate receptor 3 (IP3R3), β-tubulin and mitochondrial calcium uniport (MCU) were used as markers of endoplasmic reticulum, cytosol and mitochondria, respectively.

To clarify the membrane topology of the P2X7 subunit, this highly purified mitochondrial fraction from HEK293-P2X7R (**Figure 4A,C,E**) and N13 WT (**Figure 4B,D,F**) cells was treated with proteinase K in iso-osmotic or hypo-osmotic medium and stained with antibodies raised against the C- or N-terminal tail, or against the P2X7R extracellular loop. Proteinase K treatment in iso-osmotic medium abrogates staining with the anti-extracellular loop antibody as well as TOM20 staining, a transporter localized on the outer mitochondrial membrane, but does not affect P2X7R staining with the anti-C-terminal and the anti-N-terminal antibodies, or with an antibody against TIM23, a transporter localized on the inner mitochondrial membrane. Treatment with proteinase K in hypo-osmotic medium, which causes mitochondrial swelling and thus permeabilizes the outer membrane, decreases but does not abrogate staining with either the N- or the C-terminal antibody. TIM23 labelling is also strongly decreased in hypo-osmotic medium. Finally, membrane solubilization by Triton X-100 treatment, which makes all mitochondrial compartments accessible to proteinase K, abolishes reactivity to all antibodies. These data show that the bulky middle P2X7R domain (the extracellular loop) is exposed on the outer mitochondrial membrane, facing the cytoplasm. However, both the N- and C-termini are largely protected against proteolytic cleavage under hypo-osmotic conditions, which suggests that they might be embedded into the phospholipid bilayer. The P2X7R C-terminus is known to be highly palmitoylated (Gonnord et al. 2009) (Karasawa et al. 2017), a modification that facilitates insertion into the lipid bilayer.

**Figure 4.**
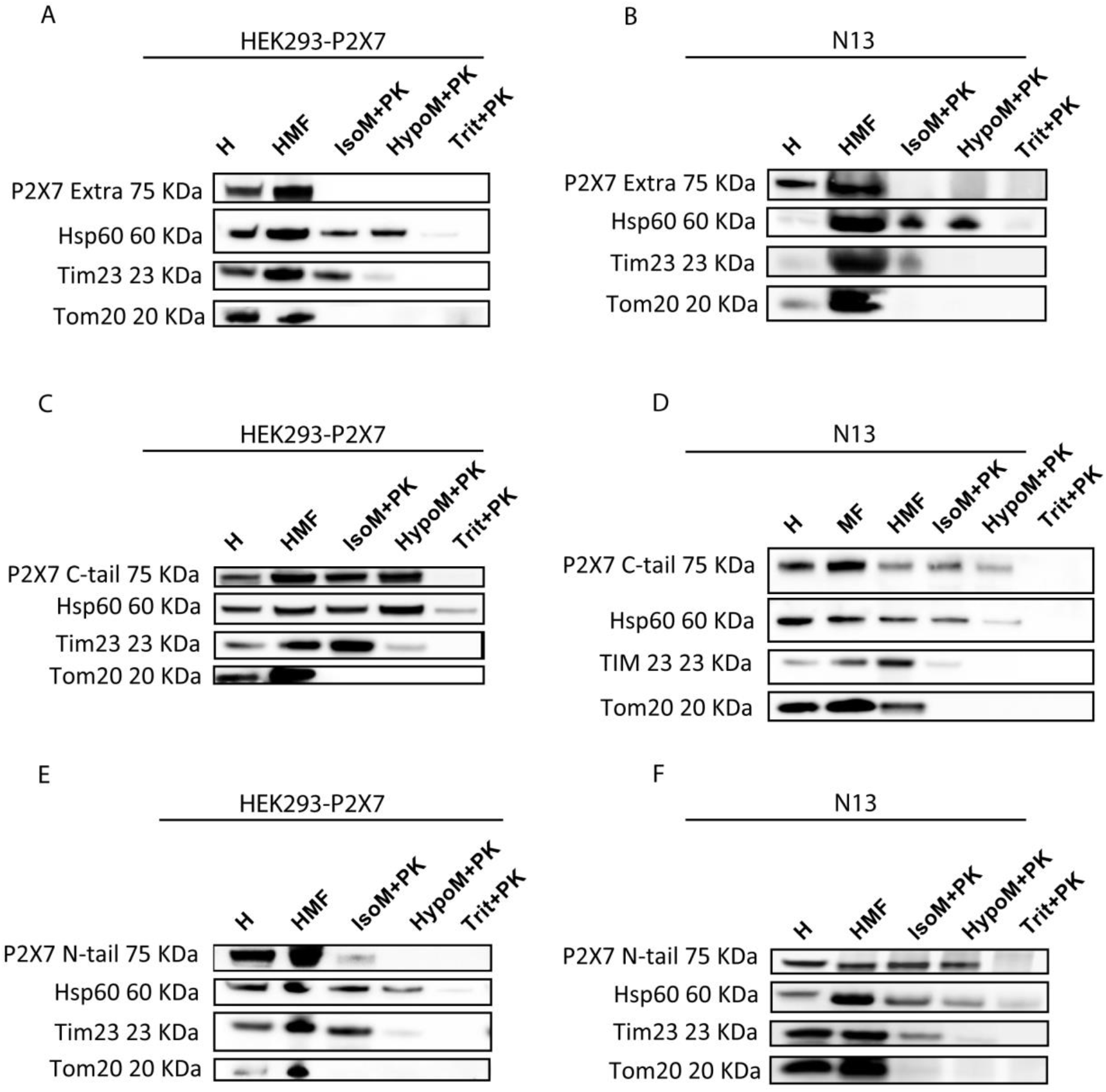
The P2X7R localizes to the outer mitochondrial membrane. Western blot analysis of a highly purified mitochondrial fraction from HEK293-P2X7R or N13 WT exposed to proteinase K (PK) (100 μg/ml) in iso- (IsoM) or hypo- (HypoM) osmotic buffer and stained with antibodies raised against the extracellular (**A**,**B**), the C-terminal (**C**,**D**) or N-terminal (**E**,**F**) domains of the P2X7R. H, whole cell lysate; MF, mitochondrial fraction; HMF, highly purified mitochondrial fraction; IsoM+PK, mitochondrial fraction incubated in iso-osmotic medium plus PK; HypoM+PK, mitochondrial fraction incubated in hypo-osmotic medium plus PK. Trit+PK, mitochondrial fraction incubated in Triton-X100-supplemented medium plus PK. Hsp60, TIM23 and TOM20 were used as markers of mitochondrial matrix, inner and outer mitochondrial membrane, respectively.

We next investigated mitochondrial P2X7R localization by confocal microscopy. To this aim, HEK293-P2X7R cells were co-labelled with an anti-P2X7R C-tail antibody and a TOM20 antibody (**Figure 5A-D**). Under resting conditions, weak P2X7R localization to the mitochondria is detected (yellow arrows in the inset in panel **A**). Mitochondrial localization is enhanced by stimulation with the P2X7R agonist benzoyl-ATP (BzATP), or with stressing agents such as rotenone or H_2_O_2_ (yellow arrows in insets in panels **B-D**, and panels **E-G**). P2X7R recruitment to the mitochondria starts soon after application of the different stimulants, and reaches a peak after about 1 h, with BzATP or rotenone, while with H_2_O_2_ peak increase is much faster (10 min), followed by a decline after 1 h, and by a delayed increase at 6 h. Total P2X7 subunit cell content does not change in response to the different stimuli (**Figure 5H**).

**Figure 5.**
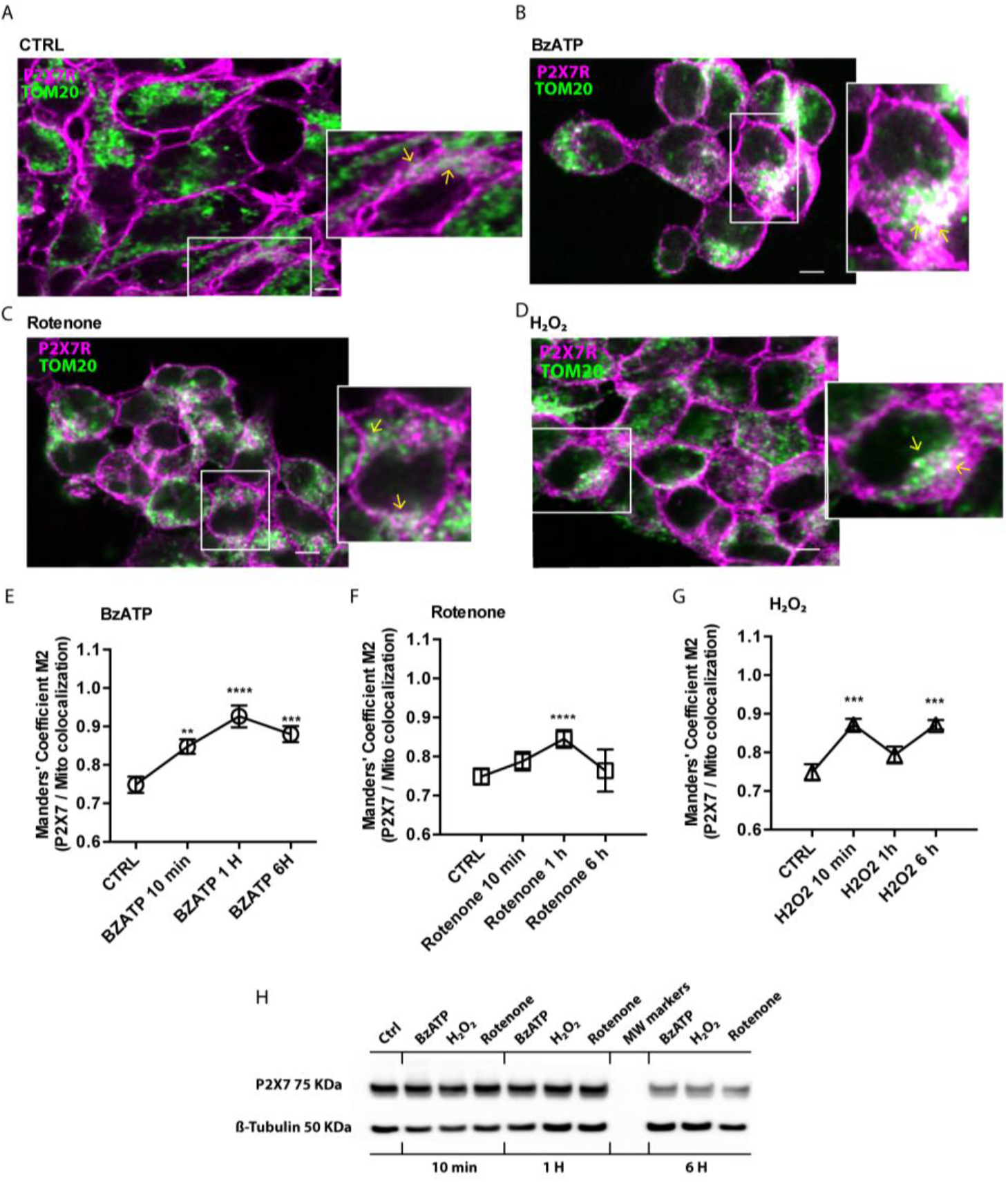
Confocal microscopy analysis of P2X7R localization. HEK293-P2X7R cell were plated on sterilized glass coverslips in 24-well plate, at a density of 5,000 cells/well in DMEM-F12 medium, and were left either untreated (**A**), or challenged with BzATP (**B**,**E**), rotenone (**C**,**F**) or H_2_O_2_ (**D**, **G**). After 1 h (BzATP or rotenone) or 10 min (H_2_O_2_) cells were fixed and stained with anti TOM20 (green) and anti P2X7R (fuchsia) antibodies. A magnification of the selected region is shown in the inset. Areas of co-localization (yellow arrow) are shown in white. Ten random fields from three independent experiments were analysed. Graphs (**E**-**G**) report quantitation of P2X7R/mitochondria localization with the Manders colocalization coefficient (expressed as percentage of P2X7R signal overlapping with TOM20 marker). Scale bar = 10 μm. Total P2X7R content at western blot under the different experimental conditions and at different time points is shown in (**H**). Exact averages ± SEM for panels **E**,**F**,**G** are shown in Supplementary Table II. P values are calculated with the two-tailed unpaired Student’s t-test. *, p < 0.05; **, p < 0.01; ***, p < 0.001.

Lack of P2X7R heavily affects several immune cell pathophysiological responses, very likely due to a defective energy metabolism (Di Virgilio et al. 2017) (Solle et al. 2001) (Hubert et al. 2010) (Karmakar et al. 2016) (Borges da Silva et al. 2018). We thus wondered whether P2X7R lack might cause effects extending beyond immunometabolism, e.g. to other tissues heavily dependent on oxidative phosphorylation. We thus investigated P2X7R localization and function in the heart.

### The P2X7R localizes to heart mitochondria and affects heart function

The P2X7R is expressed in whole mouse hearts and in a highly purified heart mitochondrial fraction (**Figure 6A**). Hearts from P2X7R-KO mice lack P2X7R immunoreactivity (**Figure 6B**), do not differ in weight from those from P2X7R-WT, but are of significantly larger size (**Figure 6C,D**). Morphometric analysis by electron microscopy revealed that mitochondria from P2X7R-KO versus P2X7R-WT mice are slightly but significantly smaller (**Figure 6E,G**), although total mass is unchanged based on the content of the TOM20 and TIM23 markers. To investigate heart function, we analyzed *in vivo* cardiac performance by High-frequency Ultrasound imaging system (Di Lascio et al. 2014) (**Figure 6H**). Left ventricle systolic volume (Vols) is slightly higher in P2X7R-KO mice (**Figure 6I**), while left ventricle diastolic volume (Vold) is lower (**Figure 6J**), but differences do not reach statistical significance. More interestingly, four basic heart indexes are significantly lower in P2X7R-KO versus WT mice: stroke volume (**Figure 6K**), fractional shortening (**Figure 6L**), ejection fraction (**Figure 6M**), and cardiac output (**Figure 6N**).

**Figure 6.**
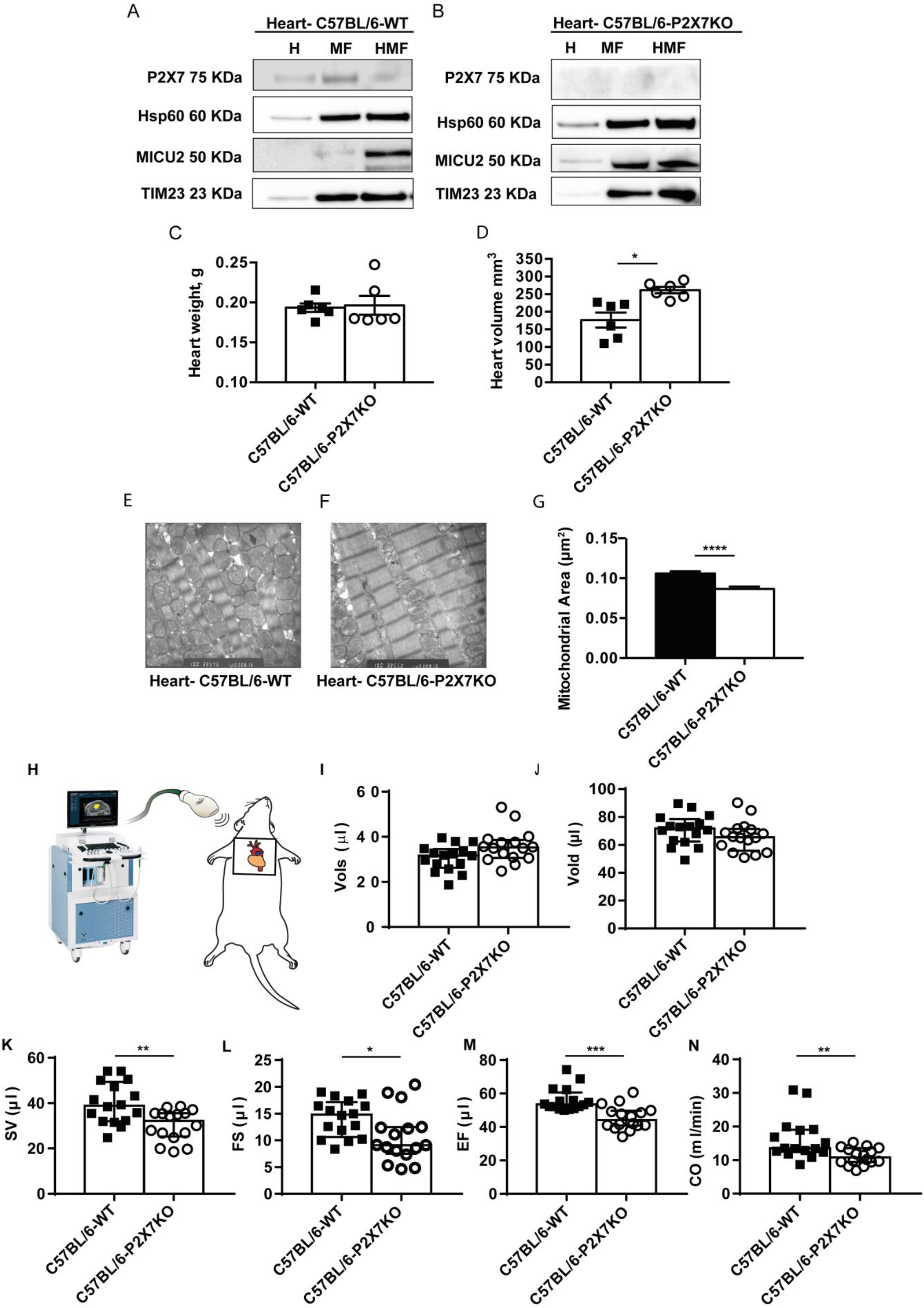
Lack of the P2X7R impairs cardiac function. Purified heart mitochondrial fractions (see Materials and Methods) from WT (**A**) or P2X7R-KO (**B**) C57BL/6 mice were analyzed for P2X7R expression by Western Blot. Hsp60, TIM23 and mitochondrial calcium uptake protein 2 (MICU2) were used as markers of mitochondrial matrix, inner or outer mitochondrial membrane, respectively. Excised whole hearts from P2X7R WT (average weight g ± SEM = 0.1935 ± 0.005343, n = 6) or P2X7R-KO (average weight g ± SEM = 0.1965 ± 0.01169, n = 6) mice were weighed (**C**), measured by caliper to assess volume (average volume mm^3^ ± SEM P2X7R WT = 176.5 ± 21.20, n = 6; average volume mm^3^ ± SEM P2X7R-KO = 261.3 ± 9.229, n = 6) (**D**), and analyzed by TEM (**E**-**G**). Mitochondrial area (μm^2^) is expressed as average ± SEM for WT (0.1059 ± 0.002917, n = 289) vs P2X7-KO (0.08657 ± 0.003239, n = 253) (**G**). In vivo heart indexes were measured with the High-frequency Ultrasound imaging system (see Methods) shown in (**H**). Vols, left ventricle systolic volume (**I**); Vold, left ventricle diastolic volume (**J**); SV, stroke volume (**K**); FS, fractional shortening (**L**); EF, ejection fraction (**M**); CO, cardiac output (**N**). Data are presented as median [IQR] of n= 16 mice for condition; *, p < 0.05; ** p, < 0.01; ***, p < 0.001; ****, p < 0.0001.

A decreased cardiac function is anticipated to have a negative impact on physical fitness, therefore we verified whether P2X7R-KO mice had a reduced performance in a Rotarod wheel test. As shown in **Figure 7A**, P2X7R-KO mice spend significantly less time on the wheel compared to P2X7R-WT. Finally we anticipated that body temperature homeostasis might be impaired in P2X7R-KO mice due to mitochondrial dysfunction. This prediction is fulfilled as average resting body temperature measured with an infrared camera is significantly lower in P2X7R-KO versus P2X7R-WT mice (**Figure 7B,C**).

**Figure 7.**
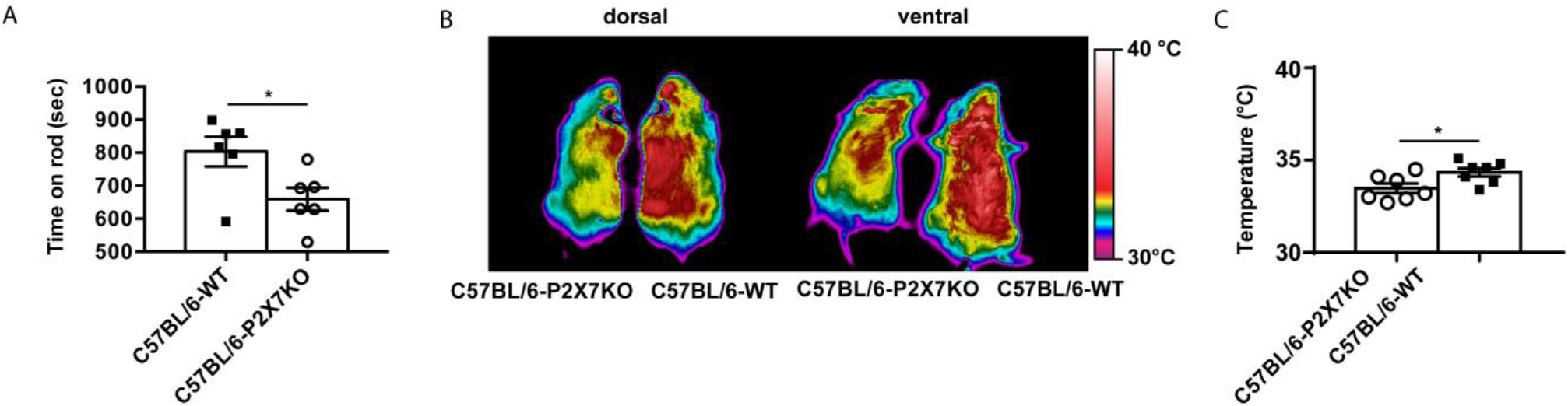
Lack of the P2X7R reduces physical fitness and surface body temperature. Physical fitness of C57Bl/6 P2X7 WT and KO mice (**A**) was investigated in a standard Rotarod test by measuring time (seconds) spent on the wheel, as described in Methods. Data are presented as average ± SEM for WT (803.3 ±44.74, n = 6) and P2X7R-KO mice (659.3 ± 34.45, n = 6). For body temperature measurement (**B**,**C**), WT and P2X7R-KO mice were placed on a warm surface in a 22° C heated room, and body images acquired and measured with a Thermacam P25 infrared Camera. Representative thermal images from one WT (right) and one P2X7R-KO (left) mouse from dorsal or ventral view (**B**). Each temperature measurement was performed in triplicate in a total of 7 mice for condition. Data are presented as average C° ± SEM for WT (33.37 ± 0.2607, n = 7) and P2X7R-KO (34.4 ± 0.226, n = 7) mice. p values are calculated with the two-tailed unpaired Student’s t-test. *, p < 0.05.

## Discussion

ATP is released into the extracellular space mainly during inflammation and muscle contraction (Bours et al. 2006) (Forrester 1966). In these conditions, extracellular ATP, acting at plasma membrane P2YRs and P2XRs, promotes immune cell recruitment, release of inflammatory cytokines, endothelial cell activation and vasodilatation (Di Virgilio et al. 2017) (Burnstock 2017). The P2X7R subtype is known for its pro-inflammatory and cytotoxic activity, but paradoxically also for its trophic effect on energy metabolism when stimulated by low agonist concentrations (Baricordi et al. 1999). While the mechanism of P2X7R-mediated cytotoxic activity has been extensively investigated (Di Virgilio et al. 1998), the molecular basis of the trophic effect is as yet poorly unknown. Several years ago we showed that tonic stimulation of the P2X7R increases the Ca^2+^ concentration of the mitochondrial matrix, supports mitochondrial metabolism and increases intracellular ATP content (Adinolfi et al. 2005). These studies complemented earlier findings again from our laboratory showing that the P2X7R, contrary to the general opinion, is not simply a cytotoxic receptor but rather a “dual function” receptor, depending on the level of activation and the metabolic state of the target cell (Baricordi et al. 1999) (Di Virgilio, Ferrari, and Adinolfi 2009). Trophic, growth-promoting activity of the P2X7R has been now demonstrated in several *in vitro* and *in vivo* systems (Di Virgilio et al. 2018) (Lara et al. 2020). Furthermore, accruing indirect evidence supports the view that the P2X7R modulates cell and whole body energy homeostasis being involved in dysmetabolisms, obesity and hormonal dysfunctions (Giacovazzo et al. 2019) (Novak and Solini 2018) (Solini and Novak 2019) (Giacovazzo, Apolloni, and Coccurello 2018). A cardio-protective activity for the P2X7R has also been proposed (Vessey, Li, and Kelley 2010).

Participation in energy homeostasis is intriguing since the only (so far) *bona fide* physiological agonist of the P2X7R is ATP, i.e. the universal fundamental intermediate in biological energy transactions. Ample evidence shows that ATP is released via non-lytic mechanisms and accumulates into the extracellular space at sites of trauma or inflammation (Di Virgilio, Sarti, and Coutinho-Silva 2020). The P2X7R itself has a leading role in setting extracellular ATP levels since it has been shown that this receptor may serve as a conduit for ATP release (Pellegatti et al. 2005) (Suadicani, Brosnan, and Scemes 2006). This is not surprising as the P2X7R macropore has a molecular cut-off of about 900 D, sufficient to allow permeation of a highly hydrophilic molecule of molecular mass of 507 such as ATP. Thus, it appears that the P2X7R has a very intricate relationship with its own agonist because on one hand it promotes ATP synthesis and transport across the plasma membrane, and on the other is a target of ATP. This intimate reciprocal relationship seems very logical in light of the pathophysiological implications of ATP accumulation into the extracellular space under a variety of conditions requiring an active energy metabolism. For example, ATP accumulates into the interstitium of exercising muscles (Forrester 1966), therefore stimulation of muscle cell P2X7R may support muscle metabolism under stress. Even more interesting in light of its role in inflammation, is P2X7R ability to “sense” ATP levels at inflammatory sites and promote a whole range of defensive responses requiring an increased energy metabolism, such as chemotaxis, phagocytosis, release of cytolytic granules, generation of reactive oxygen species (Di Virgilio et al. 2017). In this regard, the well-known low affinity of the P2X7R for ATP is of advantage because the intracellular ATP-synthetic machinery will be stimulated only when a large amount of ATP accumulates into tissue interstitium, i.e. under stress conditions when intracellular energy stores are depleted and a boost to energy synthesis is needed.

Controlled activation of the P2X7R might in principle support mitochondrial ATP synthesis in different ways, e.g. by increasing fatty acid oxidation (Giacovazzo, Apolloni, and Coccurello 2018), by facilitating glucose uptake and consumption (Glas et al. 2009), or by increasing mitochondrial Ca^2+^ levels and thus stimulating NADH oxidase activity (Griffiths and Rutter 2009). We previously hypothesized that basal activation of the P2X7R might increase basal cytoplasmic Ca^2+^ levels, and thus raise mitochondrial matrix Ca^2+^. This prediction is confirmed by experiments reported in the present work showing that P2X7R-deficient cells have a significantly lower mitochondrial Ca^2+^ concentration versus P2X7R-proficient cells (see also (Adinolfi et al. 2005)). Since mitochondrial NADH oxidase is known to be stimulated by Ca^2+^, the controlled increase in matrix Ca^2+^ is reasonably anticipated to enhance ATP synthesis (Griffiths and Rutter 2009). Thus, the P2X7R indirectly supports oxidative phosphorylation by modulating mitochondrial Ca^2+^ levels. However, NADH oxidase expression levels are also reduced in P2X7R-deficient cells. Thus, the P2X7R affects NADH oxidase in a dual fashion: by increasing its expression as well as its activity. Complex II expression is also affected by lack of P2X7R, but in opposite ways in HEK293 versus N13 R cells. In HEK293 cells, transfection with the P2X7R down-modulates Complex II expression, while in N13 R versus N13 WT cells Complex II expression is unchanged or at best slightly decreased. Down-modulation of Complex II in HEK293-P2X7R cells might be a side effect of forced P2X7R overexpression since it is not observed in cells natively expressing the P2X7R. A major role of NADH oxidase defective activity as a cause of reduced mitochondrial metabolic efficiency in P2X7R-less cells is supported by the observation that supplementation of methylsuccinate, a membrane-permeant substrate that bypasses Site I of the respiratory chain and feeds directly into Site II, restores near normal mitochondrial membrane potential.

Effect of P2X7R on NADH oxidase might be simply indirect, but based on the present data, might also be directly mediated by P2X7R mitochondrial localization, and therefore be more intimately involved in the modulation of mitochondrial physiology. Although the P2X7R is generally considered a plasma membrane channel, previous anecdotal evidence shows that it also localizes to phagosomes (Kuehnel et al. 2009), and the nuclear membrane (Atkinson et al. 2002). The physiological function of intracellular P2X7R has never been investigated, with the exception of the phagosomal localization which has been implicated in killing of phagocytosed pathogens by facilitating fusion of early phagosomes with lysosomes (Fairbairn et al. 2001). The discovery that the P2X7R is present on the mitochondria adds further complexity to the intracellular physiology of P2X7R.

Localization of the P2X7 subunit to the mitochondria in the present work was probed with three different antibodies raised against the N-terminal, the central region (extracellular loop) and the C-terminal domain of the receptor. We used both a mitochondrial fraction obtained with the standard fractionation procedure, and a highly purified mitochondrial fraction, to avoid always possible sources of contamination by intracellular organelles or the plasma membrane. Use of antibodies raised against different domains of the receptor revealed that the P2X7 subunit localizes to the outer mitochondrial membrane and allowed a tentative resolution of the membrane topology. Abolition of staining with the anti-middle domain antibody by proteinase K treatment under isosmotic conditions suggests that the middle bulky domain faces the cytoplasm, while the N- and C-termini face the inter-membrane space. Under hypo-osmotic conditions, the N- and C-termini are still partially protected from proteolysis, an effect that might be due to the tight association of these two residues with phospholipid bilayer (Gonnord et al. 2009). This membrane topology raises several intriguing questions.

The P2X7R is a cation-selective, ATP-activated channel that when expressed on the plasma membrane faces the extracellular space with the bulky, central, domain, while N- and C-termini face the cytoplasm (North and Surprenant 2000) (Karasawa and Kawate 2016). The middle domain also contains the ATP-binding sites. Therefore, ATP-binding sites in mitochondrial P2X7R face the ATP-rich cell cytoplasm. Assuming that P2X7 monomers assemble on the outer mitochondrial membrane to form a functioning P2X7R, the receptor should be constitutively activated by cytoplasmic ATP, and therefore constantly short-circuit cations across the outer mitochondrial membrane. This should not substantially alter mitochondrial energetics since it is well known that the outer mitochondrial membrane is freely permeable to small ions, but raises the question of the physiological meaning of such a pathway. On the other hand, it might well be that P2X7 subunits do not assemble into a functioning receptor, but again the meaning of these hypothetically “silent” monomers is unknown. Our data show that P2X7R expression besides improving mitochondrial metabolism also increases mitochondria size and thickness of the mitochondrial network (see also (Adinolfi et al. 2005)). These effects might be unrelated to P2X7R channel function and rather be dependent on a specific (but as yet unknown) structural activity of the isolated subunits.

Although several mechanistic details are still missing, the effect of P2X7R depletion on *in vivo* performance is dramatic. We investigated the effect of P2X7R depletion in cardiac function, as the heart is heavily dependent on mitochondrial metabolism. Cardiac mitochondria are smaller while heart volume is larger in P2X7R deficient versus proficient mice. More importantly, basic cardiac parameters (stroke volume, ejection fraction, cardiac output and fractional shortening) and physical performance (Rotarod test) are significantly lower in the absence of the P2X7R. Interestingly, surface body temperature is also lower in the P2X7R-deleted mice. Cardiac indexes in P2X7R-deleted mice are highly reminiscent of the clinical findings observed in patients affected by dilated cardiomyopathy. This is not surprising as dilated cardiomyopathy is a typical manifestation of mitochondrial disease (Meyers, Basha, and Koenig 2013).

Involvement of the P2X7R in heart dysfunction is at present very speculative. A loss-of-function mutation in the *P2RX7* gene (556C>A) has been recently found to associate with hypertrophic cardiomyopathy in a family from India (Biswas et al. 2019), but another study found no association between two *P2RX7* SNPs, the gain of function 489C>T and the loss-of-function 1513A>C, and acute heart failure in a geriatric population (Pasqualetti et al. 2017). Despite this contrasting evidence, our study highlights an as yet un-described function of the P2X7R in the regulation of mitochondrial metabolism and points to a potentially relevant role in cardiac function and physical fitness.

## Materials and Methods

### Ethical compliance

All mouse studies were performed under guidance of the University of Ferrara Institutional Animal Welfare and Use Committee and the Ethical Panel of the University of Pisa (approved protocol n. 943/2015-PR), in accordance with Italian regulatory guidelines and laws (D.Lvo 26/2014, Ministry of Health authorizations n. 75/2013-B and 76/2013-B), and the European Directive (2010/63/UE).

### Reagents

Benzoyl ATP (BzATP, cat. n. B6396), rotenone (cat. n. R8875), hydrogen peroxide solution (H_2_O_2_, cat. n. 16911) and methyl-succinate (cat n. M81101) were purchased from Sigma-Aldrich (St. Louis, MO, USA). Tetramethylrhodamine methyl ester (TMRM, cat n. T668, Molecular Probes, Leiden, The Netherlands) was dissolved in DMSO to obtain a 10 mM stock solution and then diluted in the appropriate buffer. Carbonyl cyanide *α*-[3-(2-benzothiazolyl)6-[2-[2-[bis(carboxymethyl)amino]-5-methylphenoxy]-2-oxo-2*H*-1-benzopyran-7yl]-*b*-(carboxymethyl)-tetrapotassium salt (FCCP, cat. n. C2920, Sigma-Aldrich) was solubilized in ethanol to a final stock concentration of 10 mM. For Seahorse analysis, a Seahorse XF Cell Mito Stress Test Kit was used including the following compounds: oligomycin (stock solution 100 μM), FCCP (stock solution 100 μM), and a mix of rotenone/antimycin A (stock solution 50 μM) (cat n. 103015-100, Agilent, Santa Clara, USA). Crystal violet was purchased from Sigma-Aldrich (cat n. C0775) and used as a 0.1% solution in 10% ethanol. Krebs-Ringer bicarbonate solution (KRB) contained 125 mM NaCl, 5 mM KCl, 1 mM MgSO_4_, 1 mM Na_2_HPO_4_, 5.5 mM glucose, 20 mM NaHCO_3_, 2 mM l-glutamine and 20 mM HEPES (pH 7.4), and was supplemented with 1 mM CaCl_2_.

### Cell culture and transfections

HEK293 cells were cultured in DMEM-F12 (cat n. D6421, Sigma-Aldrich) supplemented with 10% heat-inactivated fetal bovine serum (FBS, cat n. 16000044), 100 U/ml penicillin, and 100 mg/ml streptomycin (cat n. 15140130) (all from Invitrogen, San Giuliano Milanese, Italy). Stable P2X7R-transfected clones were kept in the continuous presence of 0.2 mg/ml G418 sulphate (Geneticin, cat. n. 509290, Calbiochem, La Jolla, CA). Experiments, unless otherwise indicated, were performed in the following saline solution: 125 mM NaCl, 5 mM KCl, 1 mM MgSO_4_, 1 mM NaH_2_PO_4_, 20 mM HEPES, 5.5 mM glucose, 5 mM NaHCO_3_, and 1 mM CaCl_2_ pH 7.4. N13 microglial cells, wild-type (N13 WT) and ATP-resistant (N13 R), were cultured in RPMI 1640 medium, (cat n. R0883, Sigma-Aldrich, St. Louis, MO, USA), supplemented with 10% heat-inactivated FBS, 100 U/ml penicillin, and 100 mg/ml streptomycin. Primary mouse microglia cells were isolated from 2 to 4-day-old post-natal mice as described previously (Sanz and Di Virgilio 2000). More than 98% of cells were identified as microglia using a macrophage cell-specific F4/80 rabbit mAb antibody (cat. n. MCA497, Serotec, Dusseldorf, Germany) followed by staining with Oregon Green 488 goat anti-rabbit IgG (cat. n. O-11038, Molecular Probes, Leiden, The Netherlands). Microglia cells were plated in astrocyte-conditioned high glucose-DMEM medium (cat n. 10566016), supplemented with 2 mM glutaMAX™ (Gibco Life Technologies Europe BV, Monza, Italy), 10% heat-inactivated FCS, 100 U/mL penicillin and 100 mg/mL streptomycin, and used for experiments 24 h after plating (Sanz and Di Virgilio 2000). MEFs were isolated from pregnant mice at 13 or 14 day post coitum. Mice were sacrificed by cervical dislocation, embryos were harvested, and MEFs were isolated as previously described (Missiroli et al. 2016) and cultured in DMEM medium supplemented with 10% FBS, penicillin/streptomycin, and 2 mM L-glutamine.

### Seahorse analysis

Oxygen consumption in cell lines (N13 WT, N13 R, HEK293 WT and HEK293-P2X7R) and primary cells (microglia and MEFs) was measured using the Seahorse Bioscience XF96 Extracellular Flux Analyzer (Seahorse Bioscience, Agilent, Santa Clara CA, USA). Cells were seeded in triplicate in XF96 96-well cell culture plate (Seahorse Bioscience) in a volume of 80 μl/well in DMEM complete medium at a density of 15,000/well (HEK293), 20,000/well (microglia and MEF), or 22,000/well (N13 WT and N13 R). The XF96 sensor cartridge was hydrated with 200 μl/well of XF Calibrant buffer and placed overnight in a 37°C incubator without CO_2_. On day two, incubation medium was replaced just before running the assay with Seahorse assay medium supplemented with glucose (10 mM) and sodium pyruvate (2 mM), pH 7.4, and equilibrated for 1 h at 37 °C in the absence of CO_2_. Oligomycin, to inhibit ATP synthesis, FCCP, to uncouple oxidative phosphorylation, and a mix of rotenone/Antimycin A, to block electron transport, were sequentially added at various times. Oxygen consumption rates (OCRs) were measured before and after injection of the different inhibitors. OCRs were normalized to cell content in each well determined by crystal violet staining.

### Measurement of mitochondrial membrane potential and Ca^2+^ concentration

Mitochondrial membrane potential (ΔΨm) was measured by confocal microscopy with TMRM (10 nM), at an emission wavelength of 570 nm. FCCP was used to collapses ΔΨm. Mitochondrial Ca^2+^ concentration was measured with the last-generation GCaMP probe targeted to the mitochondrial matrix (Chen et al. 2013). We chose the GCaMP6m version due to its high Ca^2+^ affinity (*K*_d_ of 167 nM). HEK293 and HEK293-P2X7 cells were grown on 24-mm coverslips, transfected with mtGCaMP6m-encoding plasmid and imaged with an IX-81 automated epifluorescence microscope (Olympus Italy, Segrate, Italy) equipped with a 40X oil immersion objective (numerical aperture 1.35) and an ORCA-R2 charge-coupled device camera (Hamamatsu Photonics, Hamamatsu, Japan). Excitation wavelengths were 494/406 nm and emission 510 nm. The 496/406 ratio is proportional to the Ca^2+^ concentration and independent of probe expression level. Analysis was performed with ImageJ.

### NADH measurement

NADH autofluorescence was measured using an inverted epifluorescence microscope equipped with a 40X oil-fluorite objective. Excitation at a wavelength of 360 nm was provided by a xenon arc lamp, with the beam passing through a monochromator (Cairn Research, Graveney Road, Faversham, UK). Emitted light was reflected through a 455 nm long-pass filter to a cooled Retiga QImaging CCD camera (Cairn Research) and digitized to 12 bit resolution. Imaging data were collected and analyzed with ImageJ. Alternatively, NADH levels in N13 WT and N13 R cells were measured using a NAD/NADH assay kit (NAD/NADH Assay Kit, Colorimetric, cat n. ab65348, Abcam, San Francisco, CA, USA), according to manufacturer’s indications.

### Subcellular Fractionation

Cells were harvested, rinsed in PBS by centrifugation at 500 g for 5 min, re-suspended in homogenization buffer (225 mM mannitol, 75 mM sucrose, 30 mM Tris-HCl, 0.1 mM EGTA, and phenylmethylsulphonyl fluoride, PMSF, pH 7.4) and gently disrupted by dounce homogenization. The homogenate was centrifuged twice at 600 X g for 5min to remove nuclei and unbroken cells, and the supernatant centrifuged at 10,300 X g for 10 min to pellet crude mitochondria. The supernatant was further centrifuged at 100,000 X g for 90 min in a 70-Ti rotor (Beckman Coulter, Indianapolis, IN, USA) at 4 °C to pellet the endoplasmic reticulum (ER) fraction. The crude mitochondrial fraction, re-suspended in isolation buffer (250 mM mannitol, 5 mM HEPES, 0.5 mM EGTA, pH 7.4), was centrifuged through Percoll gradient (Percoll medium: 225 mM mannitol, 25 mM HEPES pH 7.4, 1 mM EGTA and 30% v/v Percoll, cat n. GE17-0891-01, Sigma-Aldrich) in a 10-ml polycarbonate ultracentrifuge tube. After centrifugation at 95,000 X g for 30 min a dense band containing purified mitochondria was recovered near the bottom of the gradient and further processed as described (Wieckowski et al. 2009). Purity of the preparation was assessed by Western blot analysis of ER and mitochondrial markers. For mitochondrial fractionation, a total of at least 600 mg of isolated mitochondria was resuspended in different buffers, i.e. isosmotic buffer (10 mM Tris-MOPS, 0.5 mM EGTA-Tris, 0.2 M Sucrose, pH 7.4) with or without proteinase K (100 μg/ml), hyposmotic buffer (10 mM Tris-MOPS, 0.5 mM EGTA-Tris pH 7.4) plus proteinase K (100 μg/ml) and hyposmotic buffer plus proteinase K (100 μg/ml) and Triton (0.01% v/v Triton X-100). Samples were incubated on ice for 30 min with gentle mixing. After this time, 2 mM PMSF was added and the samples were incubated for an additional 5 min at 4°C. Proteins were precipitated with TCA/Acetone, 10% trichloroacetic acid in acetone (w/v) in a 1:6 ratio, precipitated at 4 °C for 30 min, and centrifuged at 14,000 X g for 30 min at 4 °C. Supernatant was discarded and the pellet was washed with 500 μl ice-cold acetone. Samples were centrifuge at 14,000 g for 15 min at 4 °C, resuspended in Laemmli sample buffer, and separated by SDS-PAGE.

### Isolation of mitochondria from mouse heart

Mouse hearts were washed twice with ice-cold PBS buffer to remove blood, minced, and finally resuspended in buffer A (180 mM KCl, 10 mM EDTA, 20 mM Tris HCl pH 7.4), plus 0.5 mg/ml trypsin. Samples were then incubated for 30 min at 4° C under magnetic stirring. At the end of this incubation, they were re-suspended in buffer B (180 mM KCl, 20 mM Tris HCl and 0.05 mg/ml albumin, pH 7.4). The homogenate was centrifuged twice at 4 °C for 2 min at 600 X *g*, the supernatant collected and further centrifuged at 4 °C for 10 min at 25,000 X *g*. The pellet (crude mitochondria) was re-suspended in isolation buffer and processed as describe in “Subcellular fractionation” to obtain pure mitochondria.

### Calculation of heart volume and weight

Hearts from 8-week-old C57BL/6-WT or C57BL/6-P2X7 KO mice were isolated, washed in cold PBS, weighted and measured with a caliper. The following formula was used to calculate volume: volume = π/6 × length × width^2^

### Histology

Samples were reduced and fixed in 2.5% glutaraldehyde in 0.1 M phosphate buffer, pH 7.4, and post-fixed in 2% osmium tetroxide in the same buffer, dehydrated by increasing passages in acetone and included in Araldite Durcupan ACM (Fluka address). Semi-thin sections were prepared with a Reichert Ultracut S ultramicrotome (Leica Microsystems, Buccinasco, Italy), stained with a 1% aqueous solution of toluidine blue, and observed with a light Nikon Eclipse E800 microscope (Nikon Corporation, Tokyo, Japan). Ultrathin sections were prepared with an ultramicrotome the Reichert Ultracut S ultramicrotome, counterstained with uranyl acetate in saturated solution and lead citrate (Reynolds 1963), and observed with a transmission electron microscope Hitachi H800 at 100 Kv (Hitachi High Technologies Corporation, Brughiero, Italy).

### Measurement of reactive oxygen species

The fluorogenic substrate 2’,7’-dichlorofluorescein diacetate was used to measure reactive oxygen species (ROS). Fluorescence intensity was measured with an image-based cytometer (Tali™ Image-based Cytometer, Invitrogen). Cells were pipetted into a Tali Cellular Analysis Slide and loaded into the cytometer. Bright field and green fluorescence images were captured and analysed with specific assay algorithms. Histograms are then generated to display cell size and fluorescence intensity.

### Western blot

Total cell lysates were prepared in RIPA buffer, 50 mM Tris-HCl pH 7.8, 150 mM NaCl, 1% IGEPAL CA-630 (Sigma-Aldrich), 0.5% sodium deoxycholate, 0.1% SDS, 1 mM dithiothreitol, supplemented with protease and phosphatase inhibitors (Inhibitor Cocktail cat. n. P8340, Sigma-Aldrich). Protein concentration was quantified with the Bradford assay (cat n. 5000001, Bio-Rad Laboratories, Segrate, Italy). Proteins, 15 μg/lane, were separated by SDS-PAGE, transferred to nitrocellulose membranes, and probed with the different antibodies used at the following dilutions: anti-P2X7 C-terminal, 1:500 (cat. n. APR-004, Alomone, Jerusalem, Israel); anti-P2X7 N-terminal 1:500 (cat. n. SAB2501287, Sigma-Aldrich); anti-P2X7 extracellular loop 1:200 (cat. n. P9122, Sigma-Aldrich) anti-TOM 20, 1:1000 (cat. n. D8T4N, Cell Signalling, Leiden, The Netherlands); anti-Hsp60, 1:1000 (cat.n. sc-376240, Santa Cruz); anti-β-Tubulin, 1:1000 (cat.n. T5201, Sigma-Aldrich); anti-myosin IIB 1:1000 (cat.n. M7939, Sigma-Aldrich); anti-TIM 23, 1:2000 (cat. n. ab230253 Abcam); anti-actin, 1:5000 (cat. n. A5441, Sigma-Aldrich). Densitometric analysis was performed with the ImageJ software.

### Immunofluorescence

HEK293-P2X7R cells were fixed in 4% paraformaldehyde in PBS for 15 min, washed three times with PBS, permeabilized for 10 min with 0.1% Triton X-100 in PBS and blocked in 2% BSA-containing PBS for 20 min. Cells were then incubated overnight at 4°C in a wet chamber with the following antibodies: anti-P2X7 (cat. n. P8232, Sigma-Aldrich) and anti-Tom 20 (cat n. 43406, Cell Signalling) diluted 1:100 in 2% BSA-containing PBS. Staining was then carried out with anti-rabbit Alexa 546 (Thermo Fisher Scientific Italy, Monza, Italy, cat n. A-11010) for P2X7 receptor, or with anti-mouse Alexa 633 for TOM 20 (cat n. A-21052). Cells were then washed three times with 0.1% Triton X-100 in PBS. Samples were mounted in ProLong Gold antifade (Invitrogen) and images captured with a confocal microscope (LSM 510, Carl Zeiss, Arese, Italy).

### Wound healing assay

N13 WT and N13 R cells were grown to confluence in 24-well plates, RPMI medium was replaced with low serum RPMI medium to slow down proliferation, and the wounds were made simultaneously in all wells with a sterilized pipette tip. Phase contrast pictures were taken at 0 and 24 hours. Cell migration was observed in control condition and after treatment with 2.5 mM methyl succinate. The open wound area at time 0 hours was set at 100%. Images were analysed with an open source ImageJ software.

### Infrared Thermal Imaging

Infrared thermal imaging was performed using a thermo-electrically cooled Thermacam P25 camera (Flir Systems Inc., Wilsonville, OR, USA) equipped with a scanner and 24° X 18° lens which detects a 7.5-13 μM spectral response, as previously reported (Rossato et al. 2014), and an internal calibration system with an accuracy of 0.04 °C. The focal distance was 30 cm. Images were captured and analyzed using the Flir quick report software according to manufacturer’s specifications. Each mouse (anesthetized with 2.5% isofluorane) was placed on a table at a fixed distance away the camera and images were captured in triplicate.

### High-frequency Ultrasound examination of mouse heart function

Sixteen WT and sixteen P2X7-KO male mice were examined with a high-resolution US imaging system (Vevo 2100, FUJIFILM VisualSonics Inc, Toronto, Canada) at 11-12 weeks of age. Animals were anesthetized with isofluorane in an induction chamber connected with a scavenger canister. After induction, each mouse was placed on a temperature-controlled board and the four limbs were coated with conductive paste (Signa Cream, Parker Laboratories Inc, Fairfield, Connecticut, USA) and taped with ECG electrodes. A nose cone was used to keep mice under gaseous anesthesia (1.5% isoflurane in 1 L/min of pure oxygen) during examination, and heart rate (HR), respiration rate (RR) and body temperature (T) were monitored and acquired using the Advancing Physiological Monitoring Unit attached to imaging station. The abdomen was shaved with depilatory cream (Nair, Church & Dwight Canada Corp., Mississauga, ON, Canada) and coated with acoustic coupling gel (SonoSite Cogel, Comedical Sas, Trento, Italy). For acquisition, a 40 MHz US probe (MS550D, FUJIFILM VisualSonics) held in position by a mechanical arm was employed. Images of the heart were acquired using the B-mode modality in parasternal long axis (PLAX) view, and then analysed offline to assess cardiac structure and function (VevoLAB, FUJIFILM VisualSonics Inc, Toronto, Canada). Systolic and diastolic volumes (Vols and Vold), stroke volume (SV), ejection fraction (EF), longitudinal fractional shortening (FS) and cardiac output (CO) were evaluated using the LV Trace analysis tool (Di Lascio et al. 2014) (Gao et al. 2011).

### Rotarod

The rotarod test provides information on motor parameters such as coordination, gait, balance, motivation to run and muscle tone (Rozas and Labandeira Garcia 1997). The fixed-speed rotarod test was performed as previously described (Viaro et al. 2008). Animals (six C57BL/6 WT and six C57BL/6-P2X7KO) were placed on a rotating cylinder of 8 cm in a stepwise mode at increasing speeds (from 5 to 45 rpm; 180 s each) and latency to fall was recorded. Three consecutive sessions were run. Prior to testing, animals were trained for 3 consecutive days. The cut off time was set to 30 min and the total time spent on the rod was calculated.

### Statistical analyses of biochemical and Rotarod data

All data are shown as mean ± SEM. Statistical significance was calculated assuming equal SD and variance, with a two-tailed Student’s t-test performed with the GraphPad Prism software. A p-value ≤ 0.05 was considered significant. Exact p values for each experiment were calculated and reported in the figure legends.

### Statistical analysis of cardiac parameters

The sample size was fixed taking CO as test parameter to allow for higher variability due to combination of geometrical measurements and HR assessments. Considering a standard deviation for CO measure of about 25% and a difference between wt and P2X7 KO mice of about 30% (power≥80% and alpha equal to 0.05, test significant for p<0.05), a sample size of 10 was obtained. All data are presented as median and interquartile range [25-75th IQR]. Two-sided Mann-Whitney test for independent samples was used to highlight differences between WT and P2X7 KO animals. Tests were considered statistically significant with p<0.05.

## Acknowledgments

FDV was supported by the Italian Association for Cancer Research (AIRC) grants n. IG 13025, IG 18581, IG 22883; the Ministry of Education of Italy, PRIN 2017 grant n. YTNWC, and funds from the University of Ferrara. PP was supported by AIRC, grant IG-23670; Telethon, grant GGP11139B; the Ministry of Education of Italy, PRIN 2017 grant n. E5L5P3. CG was supported by grant AIRC IG-19803; the Italian Ministry of Health, grant GR-2013-02356747; by a Fondazione Cariplo grant; by the European Research Council, ERC, 853057—InflaPML; the Ministry of Education of Italy, PRIN 2017 grant n. 7E9EPY.

## Author contribution

ACS participated in study design, performed most of the experiments, analysed data and revised the manuscript. VVP, SF, SM, ALG, MB participated in most of the experiments and revised the manuscript. PB was responsible for electron microscopy. FF, NDL and CK performed mouse ultrasound examination of mice hearts. AS participated in data analysis and interpretation, and revised the manuscript. SN and MM performed Rotarod tests and revised the manuscript. MR performed infrared thermal imaging. MRW participated in Seahorse analysis and helped in fractionation studies. CG and PP participated in study design and in data analysis, and revised the manuscript. FDV conceived the study, designed most of the experiments, analysed data and wrote the manuscript.

## Data availability

All data generated or analysed during this study are included in this published article and its supplementary information files, or are available from the corresponding author upon reasonable request.

---

**Supplementary Table I.**
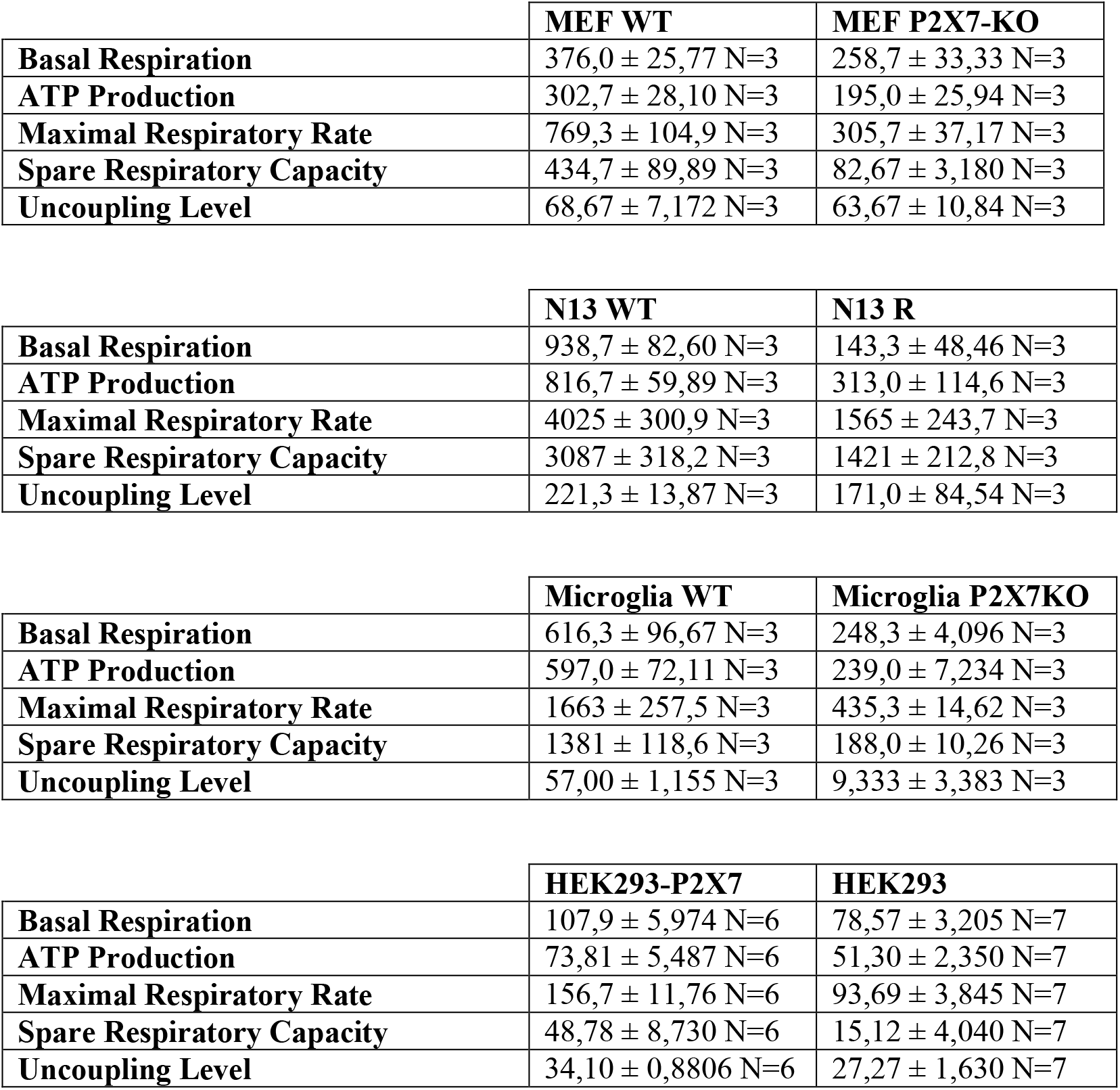

**Supplementary Table II.**

|  | **CTRL** | **BzATP** |
| --- | --- | --- |
| CTRL vs. BZATP 10 min | 0,7487 ± 0,02080 N=7 | 0,8478 ± 0,01876 N=10 |
| CTRL vs. BZATP 1 H | 0,7487 ± 0,02080 N=7 | 0,9293 ± 0,02230 N=8 |
| CTRL vs. BZATP 6H | 0,7487 ± 0,02080 N=7 | 0,8801 ± 0,02092 N=11 |

|  | **CTRL** | **Rotenone** |
| --- | --- | --- |
| CTRL vs. Rotenone 10 min | 0,7487 ± 0,02080 N=7 | 0,7877 ± 0,02371 N=7 |
| CTRL vs. Rotenone 1 H | 0,7487 ± 0,02080 N=7 | 0,8836 ± 0,008011 N=7 |
| CTRL vs. Rotenone 6H | 0,7487 ± 0,02080 N=7 | 0,7639 ± 0,05414 N=7 |

|  | **CTRL** | **H<sub>2</sub>O<sub>2</sub>** |
| --- | --- | --- |
| CTRL vs. H<sub>2</sub>O<sub>2</sub> 10 min | 0,7487 ± 0,02080 N=7 | 0,8720 ± 0,01481 N=8 |
| CTRL vs. H<sub>2</sub>O<sub>2</sub> 1 H | 0,7487 ± 0,02080 N=7 | 0,7936 ± 0,02162 N=7 |
| CTRL vs. H<sub>2</sub>O<sub>2</sub> 6H | 0,7487 ± 0,02080 N=7 | 0,8716 ± 0,01225 N=7 |

**Figure 1-Figure supplement 1.**
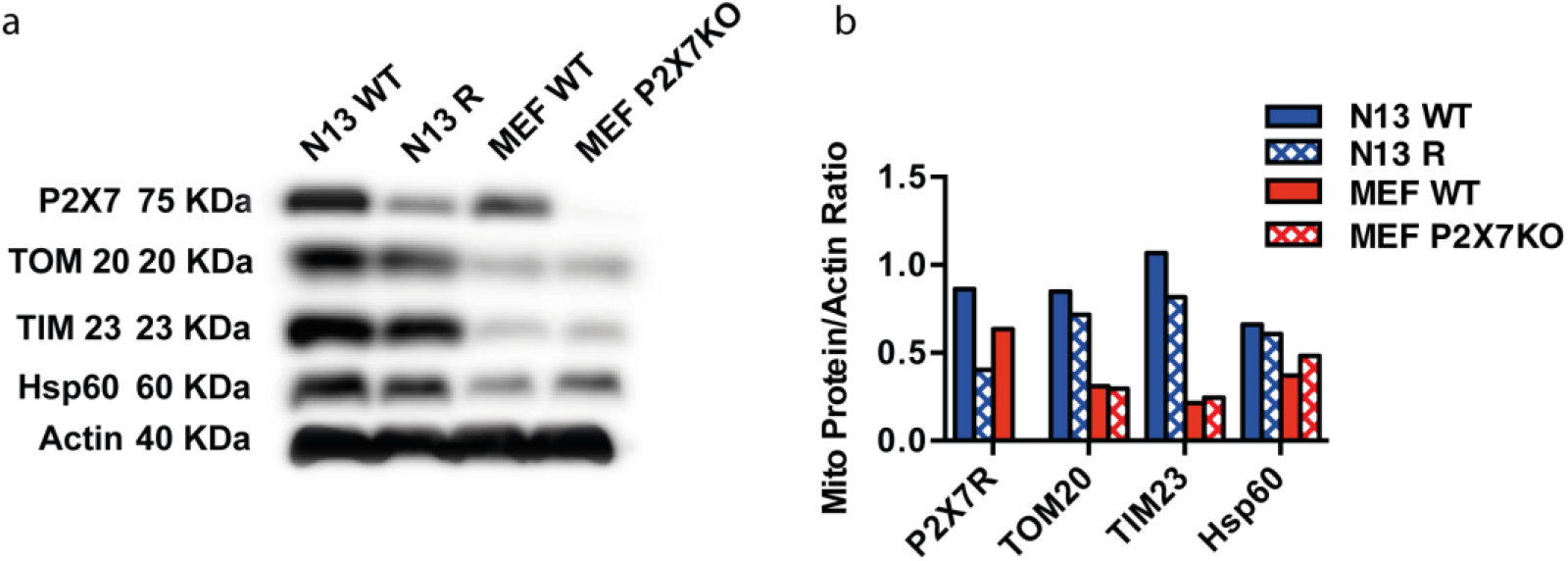
Lack of the P2X7R doesn’t affect mitochondrial cell content. Western blot analysis (**a**) and densitometry (**b**) of TOM20, TIM23 and Hsp60 content of N13 WT or N13 R, and MEF WT or MEF P2X7-KO cells.

