## Supplementary figures and images for "The P2X7 receptor localizes to the mitochondria, modulates mitochondrial energy metabolism and enhances physical performance"

### Supplemental Figure

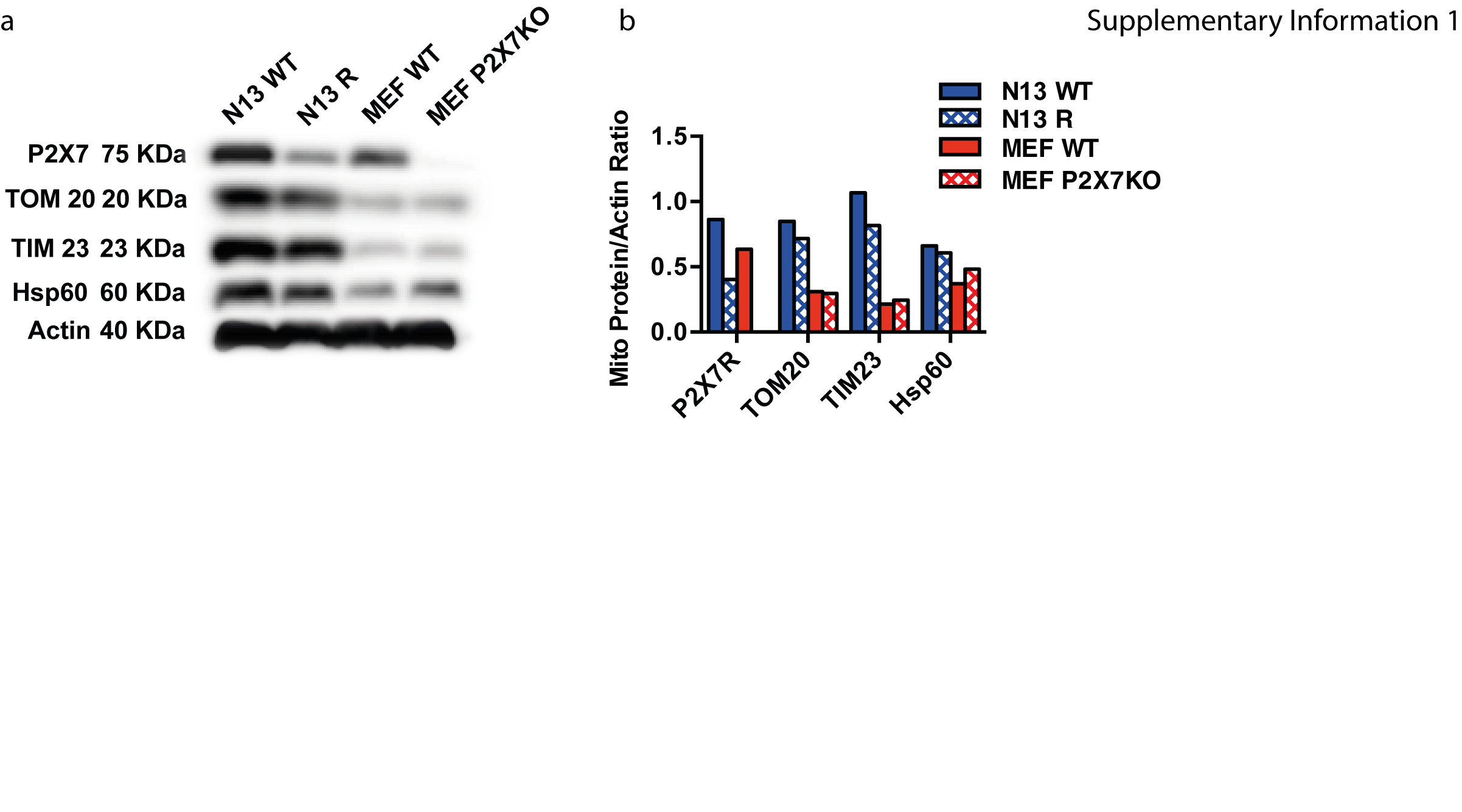
