## Supplementary Tables for "The P2X7 receptor localizes to the mitochondria, modulates mitochondrial energy metabolism and enhances physical performance"

### Supplementary Table I

| | MEF WT | MEF P2X7-KO | Unpaired t test |
|---|---|---|---|
| **Basal Respiration** | 376,0 ± 25,77 N=3 | 258,7 ± 33,33 N=3 | P value= 0,0496 d.f= 4 |
| **ATP Production** | 302,7 ± 28,10 N=3 | 195,0 ± 25,94 N=3 | P value= 0,0481 d.f= 4 |
| **Maximal Respiratory Rate** | 769,3 ± 104,9 N=3 | 305,7 ± 37,17 N=3 | P value= 0,0141 d.f= 4 |
| **Spare Respiratory Capacity** | 434,7 ± 89,89 N=3 | 82,67 ± 3,180 N=3 | P value= 0,0173 d.f= 4 |
| **Uncoupling Level** | 68,67 ± 7,172 N=3 | 63,67 ± 10,84 N=3 | n.s |

| | N13 WT | N13 R | Unpaired t test |
|---|---|---|---|
| **Basal Respiration** | 938,7 ± 82,60 N=3 | 143,3 ± 48,46 N=3 | P value= 0,0011 d.f= 4 |
| **ATP Production** | 816,7 ± 59,89 N=3 | 313,0 ± 114,6 N=3 | P value= 0,0176 d.f= 4 |
| **Maximal Respiratory Rate** | 4025 ± 300,9 N=3 | 1565 ± 243,7 N=3 | P value= 0,0031 d.f= 4 |
| **Spare Respiratory Capacity** | 3087 ± 318,2 N=3 | 1421 ± 212,8 N=3 | P value= 0,0121 d.f= 4 |
| **Uncoupling Level** | 221,3 ± 13,87 N=3 | 171,0 ± 84,54 N=3 | n.s |

| | Microglia WT | Microglia P2X7KO | Unpaired t test |
|---|---|---|---|
| **Basal Respiration** | 616,3 ± 96,67 N=3 | 248,3 ± 4,096 N=3 | P value= 0.0191 d.f= 4 |
| **ATP Production** | 597,0 ± 72,11 N=3 | 239,0 ± 7,234 N=3 | P value= 0,0078 d.f= 4 |
| **Maximal Respiratory Rate** | 1663 ± 257,5 N=3 | 435,3 ± 14,62 N=3 | P value= 0,0089 d.f= 4 |
| **Spare Respiratory Capacity** | 1381 ± 118,6 N=3 | 188,0 ± 10,26 N=3 | P value= 0.0006 d.f= 4 |
| **Uncoupling Level** | 57,00 ± 1,155 N=3 | 9,333 ± 3,383 N=3 | P value=0.0002 d.f=4 |

| | HEK293-P2X7 | HEK293 | Unpaired t test |
|---|---|---|---|
| **Basal Respiration** | 107,9 ± 5,974 N=6 | 78,57 ± 3,205 N=7 | P value=0,0009 d.f=11 |
| **ATP Production** | 73,81 ± 5,487 N=6 | 51,30 ± 2,350 N=7 | P value= 0,0021 d.f= 11 |
| **Maximal Respiratory Rate** | 156,7 ± 11,76 N=6 | 93,69 ± 3,845 N=7 | P value= 0,0002 d.f= 11 |
| **Spare Respiratory Capacity** | 48,78 ± 8,730 N=6 | 15,12 ± 4,040 N=7 | P value= 0,0036 d.f= 11 |
| **Uncoupling Level** | 34,10 ± 0,8806 N=6 | 27,27 ± 1,630 N=7 | P value=0,0049 d.f=11 |

**Supplementary Table II**

| | **CTRL** | **BzATP** | **Unpaired t test** |
|---|---|---|---|
| CTRL vs. BZATP 10 min | 0,7487 ± 0,02080 N=7 | 0,8478 ± 0,01876 N=10 | P value = 0,0033 d.f=15 |
| CTRL vs. BZATP 1 H | 0,7487 ± 0,02080 N=7 | 0,9293 ± 0,02230 N=8 | P value = 0,0001 d.f= 13 |
| CTRL vs. BZATP 6H | 0,7487 ± 0,02080 N=7 | 0,8801 ± 0,02092 N=11 | P value = 0,0006 d.f= 16 |

| | **CTRL** | **Rotenone** | **Unpaired t test** |
|---|---|---|---|
| CTRL vs. Rotenone 10 min | 0,7487 ± 0,02080 N=7 | 0,7877 ± 0,02371 N=7 | n.s |
| CTRL vs. Rotenone 1 H | 0,7487 ± 0,02080 N=7 | 0,8836 ± 0,008011 N=7 | P value = 0,0001 d.f= 12 |
| CTRL vs. Rotenone 6H | 0,7487 ± 0,02080 N=7 | 0,7639 ± 0,05414 N=7 | n.s |

| | **CTRL** | **$H_2O_2$** | **Unpaired t test** |
|---|---|---|---|
| CTRL vs. $H_2O_2$ 10 min | 0,7487 ± 0,02080 N=7 | 0,8720 ± 0,01481 N=8 | P value = 0,0003 d.f=13 |
| CTRL vs. $H_2O_2$ 1 H | 0,7487 ± 0,02080 N=7 | 0,7936 ± 0,02162 N=7 | n.s |
| CTRL vs. $H_2O_2$ 6H | 0,7487 ± 0,02080 N=7 | 0,8716 ± 0,01225 N=7 | P value = 0,0003 d.f= 12 |
